# Multiple Shifts from Peptide-Based to Metabolite-Based Venoms in Ants

**DOI:** 10.64898/2026.09.24.754088

**Authors:** Eva Vandenbroucke-Menu, Léonie Rosay, Jérôme Orivel, Alain Dejean, Corrie S. Moreau, Arnaud Billet, Vincent Courdavault, Caroline Birer Williams, Axel Touchard

**Affiliations:** UR 2106 Biomolécules et Biotechnologies Végétales, Faculté de Pharmacie, Université de Tours, Tours, France; UMR Ecologie des forêts de Guyane – EcoFoG (AgroParisTech, CIRAD, INRAE, Université de Guyane, Université des Antilles), Campus Agronomique, BP 316, 97379 Kourou Cedex, France; Université de Toulouse, Toulouse INP, CNRS, IRD, CRBE, Toulouse, France; Division of Invertebrate Zoology, American Museum of Natural History, New York, USA; UR-7417, Institut National Universitaire Champollion, Place de Verdun, 81012 Albi, France

**Author notes:** These authors jointly supervised this work. Corresponding authors: Caroline Birer Williams and Axel Touchard. These authors contributed equally to this work (Eva Vandenbroucke-Menu, Léonie Rosay).

**Keywords:** Ants, venom, peptides, alkaloids, evolution, stinging apparatus, toxins

## Abstract

Venom systems are key evolutionary innovations that have contributed to the ecological success of numerous animal lineages. In ants (Formicidae), venom exhibits an exceptional diversity in both chemical composition and delivery mechanisms. Here, we combine a synthesis of current knowledge on the evolution of ant venoms with new phylogenetic comparative analyses, integrating chemical, morphological, and phylogenetic data at the genus level. Comparative analyses across Formicidae support the hypothesis that peptide-rich venoms represent the ancestral condition in ants, as they are widely distributed across basal and derived lineages. Despite this ancestral state, we identify seven independent evolutionary transitions toward non-peptidic venom, occurring in major clades including Dolichoderinae, Formicinae, Stenammini, Solenopsidini, *Crematogaster*, *Pheidole*, and fungus-growing ants. Ancestral state reconstruction and tests of correlated evolution across 114 genera show that these seven shifts away from peptide-based venom were almost irreversible and tightly coupled with the loss of stinging capacity and a dietary shift away from predation. These transitions are consistently associated with profound morphological modifications of the venom apparatus, such as reduction or complete loss of the sting, or its transformation into non-injecting structures. Chemically, these systems are dominated by alkaloids, iridoids, formic acid, esters, and terpenoids. These repeated transitions reflect major ecological and evolutionary shifts. Lineages that rely less on individual predation and more on collective foraging or plant-based diets tend to exhibit reduced reliance on injectable peptide toxins. Instead, they favor metabolite-based venoms that can be co-opted from existing biochemical pathways and deployed in diverse ecological contexts. Conversely, peptide-rich venoms are retained in lineages subjected to strong selective pressures for rapid prey immobilization or defense against vertebrate predators. Overall, ants represent a unique model for understanding venom evolution, illustrating how shifts in chemistry, morphology, and ecological strategy can drive repeated and convergent innovations. Future integrative studies combining genomics, metabolomics, and functional assays will be essential to unravel the mechanisms underlying these transitions and their role in the extraordinary diversification of ants.

## I. Introduction

Venom is one of the most potent chemical secretions in nature, playing a significant role in the evolutionary success of many animal lineages. Although shared definitions and semantic distinctions between the terms venom and poison are still debated (Jenner, Casewell & Undheim, 2025b), several traits can be clearly attributed to venoms. Venoms are biologically active secretions that interfere with the physiological and biochemical processes of a target organism. These secretions are produced and stored in specialized glands and actively delivered to another organism through a specialized apparatus (sting, fangs, telson, tentacles, etc.), typically to subdue prey or deter predators (Nelsen *et al*., 2014; Arbuckle, 2017; Jared, Luiz Mailho-Fontana & Maria Antoniazzi, 2021; Jenner, Casewell & Undheim, 2025a). In contrast, poisons are passive chemical defenses that exert toxic effects when ingested, inhaled, or absorbed. Yet, many biological systems blur the boundaries between these categories, challenging their strict separation. Ants provide an excellent example of this complexity, highlighting a vast diversity of venomous adaptations, both in the diversity of toxins that compose them and in the morphology of the venom apparatus (Touchard *et al*., 2016a; Fox & Adams, 2022; Koch, Niedermeyer & Tragust, 2025).

The Hymenopteran venom apparatus derived from an ancestral ovipositor, which in Aculeata (ants, bees, and stinging wasps) was modified into a sting to inject venom (Robertson, 1968). Over evolutionary time, both morphological and chemical diversification have driven the emergence of a wide range of venom-related adaptations across lineages. The secondary loss of the stinging function occurred in several lineages, including Meliponini bees and the ant subfamilies Formicinae and Dolichoderinae (Blum & Hermann, 1978; Kugler, 1979; Radovic, 1981; Kumpanenko, Gladun & Vilhelmsen, 2019; Engel *et al*., 2023). Ant venoms exhibit remarkable biochemical diversity, comprising a wide range of bioactive compounds including peptides, proteins, alkaloids, iridoids, formic acid, and other small organic molecules (Touchard *et al*., 2016a), likely reflecting adaptation to diverse ecological niches and behavioral strategies. Venoms are primarily used to capture prey, with many species evolving specialized toxins for immobilizing or killing other arthropods. Venoms also exhibit antimicrobial properties, likely contributing to social immunity within the colony (Jouvenaz, Blum & MacConnell, 1972; Orivel *et al*., 2001; Téné *et al*., 2016). They can also serve as defense against vertebrate predators (Touchard *et al*., 2016a), while some venom components have communicative functions, acting as chemical signals for alarm, recruitment, or territorial marking (Morgan, 2009). Thus, the multifunctional nature of venom highlights its central role in ant ecology and social evolution.

This ecological diversity traces back to the origin of ants, estimated to have occurred in the middle Jurassic to early Cretaceous (168–140 Mya). These stem ants were ground-nesting solitary predators, with mandibles adapted for arthropod capture, and went extinct following the environmental changes of the “Cretaceous Terrestrial Revolution” (125–80 Mya), during which the rise of angiosperms opened new resources and ecological niches that fuelled the diversification of modern ants (Moreau *et al*., 2006; Jouault *et al*., 2024). Arboreal nesting evolved in tropical wet and dry climates, while other lineages adapted to angiosperm-dominated grasslands, savannas and deserts (Nelsen *et al*., 2023). Angiosperms sustained ants across these habitats through food bodies, extrafloral nectar, seed elaiosomes (rich in lipids and proteins), and indirectly through the honeydew produced by sap-feeding hemipterans. These sugary resources allowed many lineages to build large colonies whose workers forage collectively, with some species organizing group raids on prey too large to subdue individually, which they instead overpower by “spread-eagling,” a strategy in which venom plays little or no role in predation (Dejean *et al*., 2024, 2025). This diversification of ecological niches and foraging strategies alongside the angiosperm radiation raises the possibility that shifts in ant venom composition and use are themselves tied to specific ecological contexts, a hypothesis this review sets out to test by mapping venom chemistry and delivery systems onto ant phylogeny and ecology.

In this review, we reconstruct the evolutionary history of ant venom secretions and show that ants represent a unique evolutionary example among venomous animals, having repeatedly transitioned from ancestral peptide-based venoms toward metabolite toxins alongside changes in sting morphology and venom use. We also highlight the intricate relationships between morphology, chemistry, and ecological function. By mapping venom systems and their molecular compositions onto a genus-level phylogeny of ants, we identify several independent clades in which major shifts in venom composition occurred relative to the ancestral state (presumed to be peptide-rich venom delivered through a piercing sting). Rather than providing an exhaustive catalog of venom components, this review focuses on addressing the following evolutionary questions: Does stinging capacity coevolve with venom chemistry across ant lineages? How often have ant venoms transitioned from peptide-rich to non-peptide dominated compositions? Are there correlations between venom composition and specific ecological strategies or niches? Are shifts in venom chemistry associated with the ecological dominance and species richness of certain ant lineages?

## II. Phylogenetic patterns of ant venoms

In this review, we compiled and synthesized data on ant venom composition and the morphological features of the venom apparatus, including stinging capacity, across ant genera. These traits are mapped onto a genus-level phylogeny of Formicidae adapted from Nelsen, Ree & Moreau (2018) (**Figure 1**). We chose to work at the genus level because genera generally capture the major morphological and chemical diversity of ant venoms while remaining taxonomically tractable. Although exceptions exist, venom composition is often relatively homogeneous among congeneric species, making genera meaningful comparative units for large-scale evolutionary inference. Venom composition is defined here according to the dominant toxin class, such as peptides or alkaloids, based on both their relative abundance in the venom and their frequency among species within each genus. This classification does not exclude the presence of multiple toxin types within a single genus, as exemplified by the recent discovery of bioactive peptides in *Camponotus* venoms, which are otherwise dominated by formic acid (Koch *et al*., 2026). By integrating venom composition and stinging capacity with phylogenetic information, we identified seven putative clades that likely reflect independent transitions from an ancestral peptide-rich venom system toward distinct, derived chemical strategies. For the purposes of this review, these clades are named according to the major taxonomic groups they encompass: Dolichoderinae, Formicinae, Stenammini, *Crematogaster*, Solenopsidini, *Pheidole*, and fungus-growing ants. We first describe the ancestral peptide-based venom state, from which seven independent evolutionary transitions toward metabolite-based venoms subsequently emerged across the ant phylogeny.

**Figure 1.**
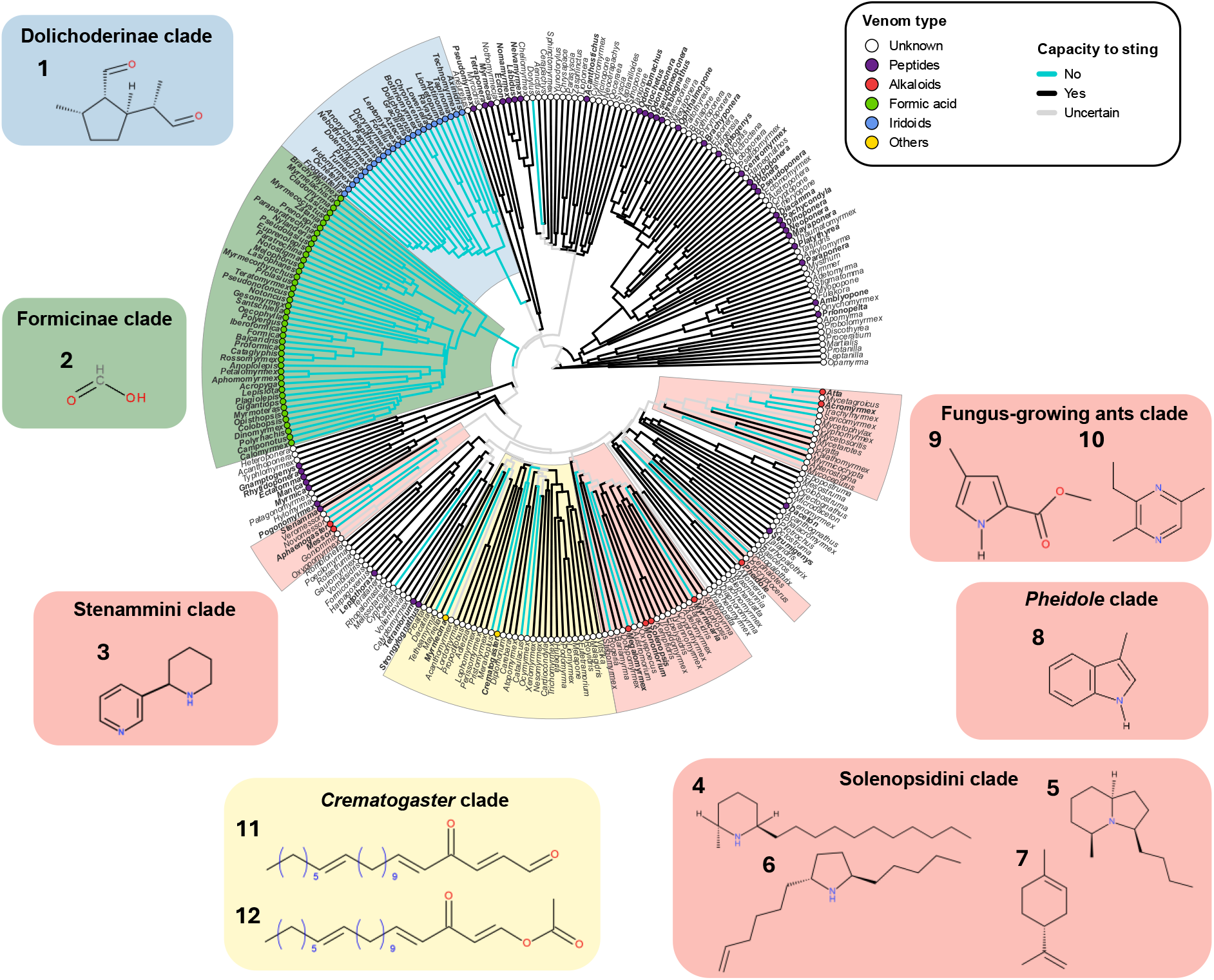
Genus-level phylogeny of Formicidae (adapted from Nelsen, Ree & Moreau, 2018) showing the distribution of venom chemical composition and stinging capacity across ant genera. Branch colour indicates the capacity to sting. Coloured circles at the tips indicate the dominant venom chemical class per genus (Peptides, Alkaloids, Formic acid, Iridoids, Others; white = unknown). Genus names are shown in bold italics where venom composition is documented. Shaded clade backgrounds highlight the seven lineages inferred as independent transitions away from the ancestral peptide-based venom state: Dolichoderinae, Formicinae, Stenammini, *Crematogaster*, Solenopsidini, *Pheidole*, and the fungus-growing ants. Chemical structures of representative metabolites are shown for each non-peptidic venom clade and are numbered to correspond to the compound numbers used throughout the manuscript: (1) iridodial; (2) formic acid; (3) anabasine; (4) solenopsin A; (5) monomorin I; (6) 2,5-dialkylpyrrolidine; (7) limonene; (8) skatole; (9) methyl-4-methylpyrrole-2-carboxylate; (10) 3-ethyl-2,5-dimethylpyrazine; (11) acetate derivatives (C19–C23); (12) aldehydes (oxidized acetates).

### (1) Peptide-based ancestral venom state

Comparative analyses of aculeate venoms, including those of Apidae, Vespidae, Pompilidae, Scoliidae and Mutillidae, consistently demonstrate that these lineages predominantly possess proteinaceous venoms dominated by small linear peptides (Konno *et al*., 2006; Chen *et al*., 2016b; Jensen *et al*., 2021; Shi *et al*., 2026). Within Formicidae, peptide-rich venoms are not confined to a single evolutionary lineage but rather are distributed across the phylogeny and mostly related to their solitary predation (Dejean *et al*., 2025). Basal ant lineages and several distantly related subfamilies possess venoms that are primarily composed of peptides with minor contributions from larger enzymatic or allergenic components, such as venom allergen 3, phospholipase A1/A2, and phospholipase B1 (Touchard *et al*., 2016a; Guido-Patiño & Plisson, 2022).

To date, peptide-based venoms have been confirmed in 41 ant genera spanning eight subfamilies: Amblyoponinae, Dorylinae, Ectatomminae, Myrmeciinae, Myrmicinae, Paraponerinae, Ponerinae, and Pseudomyrmecinae (Touchard *et al*., 2015). This extensive phylogenetic distribution supports that peptide-rich venoms are ancestral to ants, with multiple independent losses or compositional shifts occurring across lineages. Within the Myrmicinae subfamily, many lineages have transitioned away from peptide-based venoms. Nevertheless, peptide-based secretions persist in the Pogonomyrmecini and Myrmicini tribes, as well as in some Crematogastrini (e.g., *Tetramorium*) and some Attini (e.g., Dacetini) species (**Figure 1**). In contrast, some ants from the genus *Dorylus* only use venom for trail pheromone and a reduced nonfunctional sting and instead rely on hypertrophied mandibles for prey capture and defense (Billen & Gobin, 1996; A.T., personal observation). Another exception occurs within the Pseudomyrmecinae subfamily (Braekman *et al*., 1987; Merlin *et al*., 1988): an undescribed *Tetraponera* species from New Guinea produces an alkaloid-dominated venom that is delivered via a spatulate sting (Braekman *et al*., 1987; Merlin *et al*., 1988). This contrasts with the peptide-rich venoms reported for *Tetraponera aethiops* (Barassé *et al*., 2019) and *T. rufonigra* (Naephrai *et al*., 2026). Excluding these derived venom systems, we estimate that approximately 6,000 ant species (out of ≈14,000 species identified worldwide) still primarily rely on peptide-based venoms. However, substantial knowledge gaps remain, as venom composition is unknown for many genera. Recent proteo-transcriptomic studies have begun to reveal the molecular diversity of peptide-based ant venoms, providing a broader view of venom peptide repertoires for 31 species across 15 genera (Kazuma *et al*., 2017; Robinson *et al*., 2018; Touchard *et al*., 2018; Barassé *et al*., 2019; Touchard *et al*., 2020a; Aili *et al*., 2020; Barassé *et al*., 2022; Robinson *et al*., 2023b, 2024; Touchard *et al*., 2024b, 2026).

The peptide composition of ant venoms varies among species. For instance, *Rhytidoponera metallica* possesses more than one hundred distinct peptides spanning multiple functional families (Robinson *et al*., 2023b), while *Paraponera clavata* produces a venom dominated by a single peptide: poneratoxin (Aili *et al*., 2020). However, most characterized ant venoms contain approximately 10-30 peptides. Most of these belong to a single multi-gene superfamily, tentatively named aculeatoxins, which is broadly distributed across Aculeata (Robinson *et al*., 2018). Recent synteny-aware comparative genomics across 25 ant species reveal instead that venom peptides are encoded by multiple toxin gene families occupying three conserved genomic regions (GR1–GR3), which differ in their evolutionary dynamics (Weitz *et al*., 2026). Genomic investigations in two ant species (*Tetramorium bicarinatum* and *Rhytidoponera metallica*) revealed that venom genes are organized in clustered architectures showing extensive duplication and diversification, which is often associated with transposable elements (Touchard *et al*., 2024a; Isaksen *et al*., 2025). These genomic features are consistent with the rapid evolutionary turnover and adaptive diversification that are hallmarks of venom systems shaped by strong ecological interactions.

Most aculeatoxins are small, linear, amphipathic, and multifunctional peptides, many of which act directly on cellular membranes. They insert into lipid bilayers, disrupting membrane integrity and leading to cell lysis and broad cytotoxic effects across diverse taxa. These activities cause pain in vertebrates, paralysis in invertebrate prey, and antimicrobial effects (Robinson *et al*., 2018; Ascoët *et al*., 2023). In some lineages, particularly within the subfamilies Pseudomyrmecinae and Myrmeciinae, membrane-active peptides further dimerize via disulfide bonds, thereby increasing their stability and potency (Dekan *et al*., 2017; Barassé *et al*., 2019; Nixon *et al*., 2020; Touchard *et al*., 2020b; Naephrai *et al*., 2026). Beyond general membrane disruption, several ant venom peptides have evolved independently toward highly specific molecular targets. For example, neurotoxic aculeatoxins act through the modulation of voltage-gated ion channels (Robinson *et al*., 2023a; Boy *et al*., 2025). More specifically, some toxins have been shown to modulate voltage-gated sodium channels (NaVs) where they function as potent pain inducers in mammals but exhibit weak activity in insects, suggesting an adaptive role in deterring vertebrate predators rather than subduing arthropod prey (Robinson *et al*., 2023a, 2024). In addition, other venom peptides target voltage-gated potassium channels (KV and BK channels), inducing reversible paralysis on insects (Barassé *et al*., 2023; Boy *et al*., 2025). Other peptides have evolved to target G protein-coupled receptors (GPCRs). Tb1a, isolated from *Tetramorium bicarinatum*, activates the human GPCR MRGPRX2, triggering mast cell degranulation and a rapid inflammatory response (Duraisamy *et al*., 2022). *Neoponera goeldii* venom contains bradykinin-mimicking peptides that activate vertebrate BK receptors, inducing pain in vertebrate predators while remaining inactive against insect prey, toxins that have evolved independently multiple times across Hymenoptera (Shi *et al*., 2026). Additional peptide families illustrate further specialization toward particular ecological antagonists. EGF-like peptides have been identified in Myrmeciinae, including MIITX2-Mg1a from *Myrmecia gulosa*, which contains a conserved epidermal growth factor-like domain (Robinson *et al*., 2018). Mg1a mimics vertebrate growth-factor hormones, particularly those of Australian marsupials (this ant species is endemic to Australia), and induces prolonged hypersensitivity in mammalian myrmecophagous predators, suggesting evolution under pressure from specialized vertebrate antagonists (Eagles *et al*., 2022). In the neotropical ponerine *Anochetus emarginatus*, venom composition is dominated by cysteine-rich poneratoxin-like peptides encoded by a novel gene family. One of these peptides, Ae1a, induces reversible paralysis in blowflies and inhibits L-type CaV channels in human neuroblastoma cells, although only at high concentrations, with stronger effects expected in insect CaV channels (Touchard *et al*., 2016b).

The broad phylogenetic distribution of peptide-rich venoms and the repeated, independent evolution of peptides that target membranes, ion channels, G-protein-coupled receptors (GPCRs), and growth-factor signaling pathways underscore the pivotal role of predator–prey interactions in shaping ant venom evolution. The recurrent emergence of peptides that selectively target either vertebrate or invertebrate receptors provides compelling evidence that ecological pressures, particularly predation and defense, have driven the molecular diversification of ant venom peptides across stinging ant subfamilies.

### (2) The Dolichoderinae clade

The Dolichoderinae subfamily is composed of 28 genera and 725 described species (Bolton, 2026). Members of this subfamily lack a functional sting, and defensive substances are topically applied to enemies (Salado *et al*., 2023). Moreover, the venom gland is absent, and is replaced by a distinctive, large anal gland, also called the pygidial gland, a structure unique to Dolichoderinae (Blum & Hermann, 1978). This gland produces iridoids, such as iridodial **(1) (Figure 1)**, iridomyrmecin, isoiridomyrmecin, and dolichodial (Wheeler *et al*., 1977). These cyclopentanoid monoterpenoids or iridoids, named after the ant *Iridomyrmex detectus*, are well-known specialized metabolites found in plants and insects. In both groups, iridoid biosynthesis involves a complex series of enzymatic transformations, starting from the monoterpenoid precursor geranyl diphosphate (GPP) (Beran *et al*., 2019). The identification of a P450 enzyme with geraniol 8-hydroxylase activity in *Linepithema humile*, an invasive ant species originally from Argentina, which is evolutionarily distinct from the geraniol 8-hydroxylase P450 previously characterized in insects, supports the hypothesis that iridoid biosynthetic pathways evolved independently in Dolichoderinae and other insect groups (Datta *et al*., 2026). The monoterpenoid actinidine has been identified in the pygidial gland and is hypothesized to result from the reaction between ammonia and iridodial present in the gland (Cavill *et al*., 1982). In some *Dolichoderus* species, such as *Do. bispinosus*, actinidine is reported as the sole constituent (Janssen *et al*., 1995).

The amount of iridomyrmecin in the pygidial gland of *Li. humile* varies markedly among supercolonies, but this variation shows no association with invasive success or with invasive versus native ranges (Salado *et al*., 2023). *Dolichoderus scabridus* and *Iridomyrmex rufoniger* also exhibit intercolonial differences in pygidial gland secretion, with colonies producing either methylheptenone and iridodial or dolichodial as the predominant compounds (Cavill & Hinterberger, 1960). These differences may reflect the existence of cryptic species.

Pygidial gland iridoids serve primarily as defensive compounds in most Dolichoderinae species. *Linepithema humile* applies iridomyrmecin, which has antibiotic and insecticidal activity, and dolichodial directly onto the cuticle of competitors during conflicts. These compounds irritate heterospecifics but also elicit attraction of nestmates (Vander Meer, 2012). Similar compounds suppress aggression from other ant species. In *Li. humile*, iridoids also mediate recruitment or necrophoresis depending on release quantity (Welzel *et al*., 2018). Dolichodial and iridomyrmecin disappear within an hour after death, triggering necrophoretic behavior in nestmates (Choe, Millar & Rust, 2009). Iridodials from *Tapinoma simrothi* and *Ir. rufoniger* likewise induce nestmate recruitment (Cavill & Hinterberger, 1960; Van Oudenhove *et al*., 2012).

### (3) The Formicinae clade

Formicinae is a diverse and ecologically dominant ant subfamily with 55 genera and more than 3,200 described species to date (Bolton, 2026). It represents a major evolutionary shift in the ant venom system. Formicine ants are entirely stingless, relying on formic acid-based venom (Koch *et al*., 2025). However, antimicrobial peptides have recently been characterized in the venoms of *Camponotus* species, suggesting that the venom composition of formicine ants is more complex than a system solely based on formic acid (Koch *et al*., 2026). Notably, synteny-aware comparative genomics across 25 ant genomes reveals that formicine species that spray formic acid typically retain only a single copy, or in some cases no copies, of canonical venom peptide genes at major toxin loci (Weitz *et al*., 2026). The loss of the sting has been evolutionarily compensated by the development of a unique morphological adaptation for venom excretion: the acidopore. This round nozzle-like opening, located at the end of the cloaca and surrounded by a fringe of hairs, facilitates the precise spraying of venom from a distance or directly into bite wounds (Hung & Brown, 1966).

Formic acid, also known as methanoic acid (CH₂O₂), is a specialized metabolite derived from the tetrahydrofolate (THF) cycle. Serine and glycine donate methylene groups to THF via either serine hydroxymethyltransferase or the glycine cleavage system. After oxidation of 5,10-methylene THF by methylene THF dehydrogenase, 5,10-methenyl THF is hydrolyzed to 10-formyl THF by 5,10-methenyl THF cyclohydrolase. Two subsequent reactions can occur: either formate is released by 10-formyl THF synthetase, which phosphorylates ADP to ATP, or formic acid is released by 10-formyl THF hydrolase (Koch *et al*., 2025). According to Hefetz & Blum (1978), phosphorylation of ADP to ATP is thought to be a protective mechanism for excretory cells. The concentration of formic acid shows both inter-and intraspecific variation, influenced by factors such as species, colony, caste, age, morphology, and reproductive status (Melander & Brues, 1906). Additionally, it can fluctuate based on fighting behavior and application methods (Stumper, 1923a, 1923b, 1950, 1952). Although formic acid is the dominant compound of formicine venom, other compounds can be present. Among volatiles, acetic acid is sometimes present in large amounts, along with short-chain aliphatic carboxylic acids. Small amounts of aromatic compounds may also be present, and decahydroquinolines have been identified in the venom of the weaver ant *Oecophylla smaragdina* (Das *et al*., 2014; Mekonnen *et al*., 2021; Xu *et al*., 2023). The inorganic fraction contains ammonia, calcium, iron, and phosphate (Flury, 1919; Stumper, 1959, 1960), while the organic fraction includes amino acids that may contribute to venom toxicity, such as non-proteinogenic GABA, or play a role in venom dispersal (Ghent, 1961; Hermann & Blum, 1968).

Formicine ants release venom by spraying it from a distance or directly into bite wounds, where formic acid **(2) (Figure 1)** causes rapid tissue destruction (Koch *et al*., 2025). Formic acid can also be applied onto insect cuticles and then enters through the tracheal system (Löfqvist, 1977). At high concentrations, formic acid is highly corrosive, while at lower concentrations it is cytotoxic by inhibiting the cytochrome c oxidase, an essential mitochondrial enzyme (Nicholls, 1975). In addition to its defensive and offensive functions, formic acid can be involved in nest hygiene by inhibiting the proliferation of pathogenic microorganisms, detoxification of alkaloids (e.g. formylation, protonation/sequestration), herbicide, as well as in triggering alarm or nestmate recruitment (Frederickson, Greene & Gordon, 2005; Tragust *et al*., 2013a, 2013b, 2020; LeBrun, Jones & Gilbert, 2014; Tranter *et al*., 2014; Koch *et al*., 2025).

### (4) The Stenammini clade

The tribe Stenammini, within the Myrmicinae subfamily, comprises seven genera and 458 described species (Bolton, 2026). Most belong to *Aphaenogaster* and *Messor*, which differ markedly in their ecology; the former being omnivorous, whereas the latter is primarily granivorous. Stenammini ants possess a reduced venom apparatus characterized by flanges on the sting shaft, although detailed morphological examination remains scarce (Kugler, 1979). The venom consists mainly of pyridine alkaloids originally identified from tobacco plants, including anabasine **(3) (Figure 1)**, anabaseine, 2,3’-bipyridyl, and myosmine, along with pyrazine alkaloids, such as 3-ethyl-2,5-dimethylpyrazine (Fox & Adams, 2022). Mannich-like reactions are widespread in alkaloid biosynthesis, involving the attack of a carbon nucleophile on an iminium electrophile, with some steps being enzyme-catalyzed while others may occur spontaneously under specific physiological conditions. The biosynthetic pathways of pyridine alkaloids in ants remain undescribed but may parallel those in plants for which anabasine biosynthesis begin with lysine decarboxylation to cadaverine, followed by its oxidation into 5-amino-pentanal, which spontaneously cyclizes into Δ1-piperideine. Anabasine results from a final cyclization between Δ1-piperideine and nicotinic acid N-glucoside, coordinated by a cascade of four enzymes, including two oxidoreductases and a key β-glucosidase (Schwabe *et al*., 2026).

Venom composition within the Stenammini clade shows marked interspecific variation. In *Messor*, anabasine generally dominates (Leclercq *et al*., 2001; Co *et al*., 2003), whereas *Aphaenogaster* species primarily produce anabaseine (Wheeler *et al*., 1981; Attygalle *et al*., 1998; Lenoir *et al*., 2011). Nonetheless, exceptions occur. The venom of *Me. capensis* contains anabaseine as the major alkaloid and anabasine as a minor compound (Brand & Mpuru, 1993), while *Me. arenarius* produces a more complex blend of pyridine derivatives, pyrazines and piperidines (Co *et al*., 2003; Cruz-López *et al*., 2006). Some *Messor* species lack alkaloids: *Me. structor* and *Me. rugosus* possess venoms dominated by the linear alkane heptadecane, and *Me. barbarus* mainly produces methylbenzoate (Leclercq *et al*., 2001; Co *et al*., 2003). Yet, a later analysis of *Me. rugosus* reported the presence of nicotinic alkaloids (Cruz-López *et al*., 2006). In contrast, the venom of *Stenamma debile* consists primarily of peptides, although the possible presence of alkaloids has not been tested yet (Barassé *et al*., 2022). The venom chemistry of related genera, such as *Goniomma*, *Novomessor* and *Veromessor*, remains unexplored.

Pyridine alkaloids, known for their insecticidal properties, serve multiple roles in ants. In *Ap. tennesseensis* and *Ap. fulva*, anabaseine acts as a defensive compound, with the latter species also using it to attract conspecifics (Wheeler *et al*., 1981; Blum, 1992). In *Me. ebeninus*, anabasine functions as an alarm pheromone (Coll, Hefetz & Lloyd, 1987). These alkaloids may also function as trail pheromones in Stenammini ants. In *Ap. rudis*, a mixture of anabasine, anabaseine, and 2,3’-bipyridyl synergistically triggers trail-following behavior (Attygalle *et al*., 1998), while in *Me. bouvieri*, 3-ethyl-2,5-dimethylpyrazine appears to play a similar role (Jackson, Wright & Morgan, 1989).

### (5) The Solenopsidini clade

The Solenopsidini tribe, within the Myrmicinae subfamily, comprises 23 genera, and 762 described species (Bolton, 2026). Their venom is mostly characterized by containing alkaloids, and only a few peptides have been reported to date. Most research has focused on a few genera: *Solenopsis*, *Megalomyrmex*, *Monomorium* and *Myrmicaria*. Among them, *Solenopsis* is by far the best studied, with detailed investigations of solenopsin alkaloid isomers and enantiomers (Xu & Chen, 2023) and analysis of venom composition in hybrid species (Chen *et al*., 2019). The venom apparatus of Solenopsidini displays distinctive morphological modifications, including the reduction of the structure and an expansion of the distal surface of the aculeus. In certain genera, such as *Megalomyrmex*, *Monomorium*, and *Myrmicaria*, the sting is spatulate (Kugler, 1979; Kaib & Dittebrand, 1990; Kenne *et al*., 2000a), whereas in *Solenopsis*, it retains a typical morphology that allows efficient venom injection (Xu & Chen, 2023). Gaster-flagging behaviors, where ants raise and vibrate their abdomen to disperse venom droplets, have been documented in *Solenopsis*, *Megalomyrmex* and *Monomorium* (Adams *et al*., 2015; Xu & Chen, 2023).

In *Solenopsis*, venom alkaloids fall into several molecular classes, including piperidines, pyridines and pyrrolidines. Among these, the diverse group of solenopsins exemplifies mixed metabolic origin, combining a piperidine heterocycle derived from the lysine pathway, with a hydrophobic side chain from fatty acid metabolism, as seen in solenopsin A **(4) (Figure 1)** (MacConnell, Blum & Fales, 1970; Xu & Chen, 2023). Structural diversity among solenopsins results from variation in substituent configuration, chain length, and alkyl chain unsaturation (Leclercq, Braekman & Daloze, 1996). Minor proteinaceous components are also present (Jones, Blum & Fales, 1982) and venom composition is known to vary with morph, caste, age, and body size (Brand *et al*., 1972; Deslippe & Guo, 2000). In *Monomorium*, the venom contains monomorins, which belong to the pyrrolidine, piperidine and indolizidine classes of alkaloids (Jones *et al*., 1982; Fox & Adams, 2022). Monomorin I **(5) (Figure 1)**, for instance, is likely synthesized from L-lysine via cadaverine-derived intermediates, forming an indolizidine backbone through enzyme-mediated cyclization, although the enzymes involved remain unidentified. In *Megalomyrmex*, pyrrolidines, pyrrolizidines, and indolizidines predominate (Jones *et al*., 1991; Adams *et al*., 2015; Sozanski *et al*., 2020; Fox & Adams, 2022), except in *Me. mondaboroides*, whose venom is characterized by piperidines, some shared with *Solenopsis* and *Monomorium* (Adams *et al*., 2015). Transcriptomic analyses of venom glands in *Megalomyrmex milenae* also revealed toxin-like peptides, including putative pore-forming and antimicrobial components, suggesting a more complex venom profile. However, these candidates have not yet been confirmed by mass spectrometry and therefore remain to be validated (Sozanski *et al*., 2026).

Among Solenopsidini, venom fulfils multiple ecological functions, including insecticidal, repellent, antimicrobial and communicative roles. Piperidine alkaloids in *Solenopsis* and pyrrolidines and pyrrolizidines in *Megalomyrmex* act as potent insecticides (Jones *et al*., 1982; Evershed, Morgan & Cammaerts, 1982; Sozanski *et al*., 2020), while venom also serves as a repellent in *Solenopsis*, *Megalomyrmex*, and *Monomorium* (Blum *et al*., 1980; Andersen, Blum & Jones, 1991; Adams *et al*., 2015). In *Monomorium minimum*, 2,5-dialkylpyrrolidines **(6) (Figure 1)** act as repellents, allowing workers to deter competitors and to retrieve prey (Adams & Traniello, 1981; Chen *et al*., 2016a). Some venoms also act in communication, notably in *Monomorium pharaonis*, in which monomorin I, an indolizidine alkaloid, and a pyrrolidine alkaloid synergize Dufour’s gland pheromone faranal to elicit trail following (Evershed *et al*., 1982). Antimicrobial and antifungal activities have been demonstrated in *Megalomyrmex* and *Solenopsis* venoms and contribute to brood sanitation (Sozanski *et al*., 2020; Xu & Chen, 2023). Beyond these functions, venom plays a role in social parasitism and raiding behavior. In *Solenopsis fugax* and *Monomorium* species, venom acts as a repellent, facilitating brood or food theft (Andersen *et al*., 1991; Adams *et al*., 2015). In *Megalomyrmex,* venom has evolved as a pacifying agent: the venom of *Me. mondabora* and *Me. sylvestrii*, rich in pyrrolidines and pyrrolizidines, induce thanatosis (death-feigning), rather than aggression in the fungus-growing ant *Cyphomyrmex costatus* enabling coexistence within attine nests (Adams *et al*., 2000). In *Me. symnetochus*, venom acts as propaganda allomone, protecting its host *Sericomyrmex* by preventing recruitment and disrupting raids by *Gnamptogenys hartmani* (Adams *et al*., 2013).

Lastly, the genus *Myrmicaria* is characterized by a modified sting whose distal portion is membranous and blunt, a morphology that facilitates topical venom application (Kaib & Dittebrand, 1990; Kenne *et al*., 2000b). Their venom is dominated by monoterpenes, most notably limonene **(7) (Figure 1)** (Blum, 1992), together with indolizidine, pyrroloindolizidine and lehmnizidine polycyclic alkaloids collectively termed myrmicarins (Jones *et al*., 2007; Ondrus, Ümit Kaniskan & Movassaghi, 2010; Fox & Adams, 2022). Venom is primarily directed against prey and also functions as an alarm pheromone. In *My. opaciventris*, venom alkaloids exhibit paralytic effects on target organisms (Ondrus *et al*., 2010). By contrast, *My. eumenoides* produces venom comprising two functionally interacting fractions: a highly volatile fraction, consisting of 97% limonene, which acts as an alarm pheromone, and a less volatile alkaloid-rich fraction with suspected insecticidal activity. The alkaloid fraction modulates limonene evaporation prolonging its activity while limonene enhances the penetration of alkaloids across the arthropod cuticle, presumably by disrupting the lipid-rich cuticular barrier and facilitating their diffusion (Kaib & Dittebrand, 1990).

### (6) The *Pheidole* clade

This clade, previously called the Pheidolini tribe, includes the genera *Pheidole*, *Procryptocerus*, and *Cephalotes*. *Pheidole* is the most successful ant genus, encompassing 1,250 described species, and represents an ecologically significant group within the subfamily Myrmicinae (Wilson, 2003). These ants exhibit a dimorphic worker caste consisting of majors (also referred to as soldiers) and minors. They are known for their aggressive behavior and high competitiveness with other species. The genus is predominantly omnivorous, with some species specializing in seed harvesting while others capture live insects. However, they do not appear to rely on chemicals to subdue their prey (Dejean *et al*., 2007). Colony defense and food protection are typically achieved through cooperative fighting rather than stinging. Group retrieval of prey or defense against intruders is frequently performed by “spread-eagling” the target (Moreau, 2008). Despite being often described as non-venomous due to their atrophied, non-functional sting, *Pheidole* ants do possess a venom system with well-developed venom glands and reservoirs. In *Pheidole jelski*, the venom reservoir in soldiers is hypertrophied, with the venom appearing brown rather than the transparent venom found in minor workers (A.T., personal observation), suggesting a specialized defensive function. Similarly, soldiers of *Pheidole fallax* possess hypertrophied venom reservoirs that emit a fecal odor attributed to an indole compound, likely skatole **(8)** (3-methylindole) **(Figure 1)** (Law, Wilson & Mccloskey, 1965). These observations indicate that the venom may serve at least a repellent function for defensive purposes. Very few studies have analyzed the venom reservoir content in *Pheidole*. Several pyrazines have been detected in the venom reservoirs of minor workers of *Ph. pallidula* and major workers of *Ph. sinaitica* (Ali, Jackson & Morgan, 2007). These findings highlight the need for further investigation into the chemical diversity and ecological roles of *Pheidole* venoms.

In *Cephalotes*, no information is currently available regarding venom composition. These ants have lost the ability to sting and possess a reduced sting. They are primarily herbivorous, with a diet based on plant-derived resources, supplemented by pollen, bird feces, and vertebrate urine. Dissections of *Cephalotes atratus* reveal that workers retain a well-developed venom system, including filamentous venom glands and a venom reservoir filled with a yellowish, oily secretion (A.T., personal observation). This suggests that, despite the loss of a functional sting, the venom apparatus remains active. The function of this secretion remains unknown, but it may be involved in chemical communication, potentially acting as a trail pheromone. Notably, *Cephalotes umbraculatum* exhibits a defensive posture reminiscent of *Crematogaster* species, raising the abdomen over the head while releasing a whitish secretion (A.T., personal observation). This behavior suggests that venom-derived compounds may also serve as repellent or defensive functions. In contrast, no behavioral observations or chemical data are currently available for the closely related genus *Procryptocerus*, leaving the function and composition of their venom system entirely unexplored.

### (7) The fungus-growing ants clade

The fungus-growing ants clade comprises 14 genera and 207 extant species. The clade is traditionally divided into two major groups: the “lower fungus-growing ants” which comprises most genera and “higher fungus-growing ants”. All species are obligate fungus farmers and fungiculture is considered to have originated once. Lower Attini cultivate Leucocoprineae or Pterulaceae fungi from organic detritus, i.e. insect frass and dead vegetation, while higher fungus-growing ants are leaf-cutters of the genera *Acromyrmex* and *Atta*, associated to a single fungal cultivar, *Leucoagaricus gongylophorus*, incapable of free-living existence (Schultz & Brady, 2008). Ants protect the fungal cultivars from competitors and pathogens using fungicidal compounds produced by their cuticular bacterial microbiota, while the fungus provides a primary food for the colony (Currie *et al*., 1999). Consequently, fungus-growing ants have become a key model for studying coevolution and mutualism (Currie, 2001; Mueller *et al*., 2005). However, relatively little is known about their venom, as the clade is generally not considered venomous despite possessing a venom gland. In terms of the venom apparatus, *Sericomyrmex* is known to have only a vestigial sting (Adams *et al*., 2013).

Venoms of *Atta* and *Acromyrmex* species contain pyrrole and pyrazine-derived alkaloids that act as trail pheromones (Evershed & Morgan, 1983). Methyl-4-methylpyrrole-2-carboxylate **(9) (Figure 1)**, alone or combined with pyrazines, elicits trail-following behavior in several species of both genera (Tumlinson *et al*., 1971; Cross *et al*., 1979, 1982; Evershed & Morgan, 1983; De Oliveira *et al*., 1990; Do Nascimento *et al*., 1994; Campos *et al*., 2016). By contrast, trail-following in *At. sexdens sexdens* and *At. sexdens rubropilosa* is triggered by 3-ethyl-2,5-dimethylpyrazine **(10) (Figure 1)** in association with pyrroles (Cross *et al*., 1979; Evershed & Morgan, 1983). In *Trachymyrmex*, however, neither pyrazines nor pyrroles have been reported. Instead, this genus produces a mixture of alkanes (most notably undecane) from the Dufour’s gland, which serves as both alarm and aggregation pheromone (Adams *et al*., 2012).

### (8) The Crematogaster clade

This clade includes the genera *Crematogaster* and *Myrmecina*, which cluster together despite the limited and uneven knowledge of venom composition across these genera and their close relatives.

*Crematogaster* is another ecologically successful genus of ants that mostly employs a strictly defensive contact venom to ward off competing ants, termites, and other arthropods (Marlier, Quinet & de Biseau, 2004). However, an offensive role has been observed in the prey capture for *Cr. striatula*, whose venom acts at a distance (Rifflet *et al*., 2011), as well as in another undetermined African species, where it is applied topically on prey (Richard, Fabre & Dejean, 2001). However, most *Crematogaster* species appear highly generalist omnivores that rarely use venom for predation (Buren, 1958).

The genus has followed a unique evolutionary path for its venom system, characterized by spatula-shaped sting tip (Buren, 1958) and a dorsally inserted post-petiole that enables the gaster to bend over the head, a hallmark defensive posture. Venom composition, studied mainly in *Cr. scutellaris* and some other European ant species, consists of acetate derivatives with acetylated C19-C23 carbon chains **(11) (Figure 1)** stored in Dufour’s gland (Daloze *et al*., 1987, 1991). During secretion, venom gland enzymes, an esterase and an alcohol oxidase, convert these acetates into unstable, toxic aldehydes **(12) (Figure 1)**, thereby indirectly protecting Dufour’s gland cells (Pasteels, Daloze & Boeve, 1989). This reaction also releases acetic acid, which acts as an alarm pheromone. Similar compositions were found in three other unidentified *Crematogaster* species from Papua New Guinea (Leclercq *et al*., 1997), while *Cr. brevispinosa* from Brazil contains distinctive furanocembranoid diterpenes (Crematofuram) (Leclercq *et al*., 2000). Given the remarkable diversity of the genus, with over 500 species, venom chemical diversity likely extends far beyond these examples.

Another non-stinging myrmicine within this clade, *Myrmecina graminicola*, possesses a markedly hypertrophied venom reservoir containing large quantities of acetate and propionate esters (Lenoir *et al*., 2018), further supporting the idea that acetate-based venom systems have evolved convergently or been retained within this lineage.

## III. Evolutionary Analyses

To characterize how venom chemistry has evolved and whether its evolution is linked to other ecologically relevant traits, phylogenetic comparative analyses of venom chemical composition were conducted across the ant phylogeny, retaining all genera (N=114) for which venom-type information was available (**Figure 2A**).

**Figure 2.**
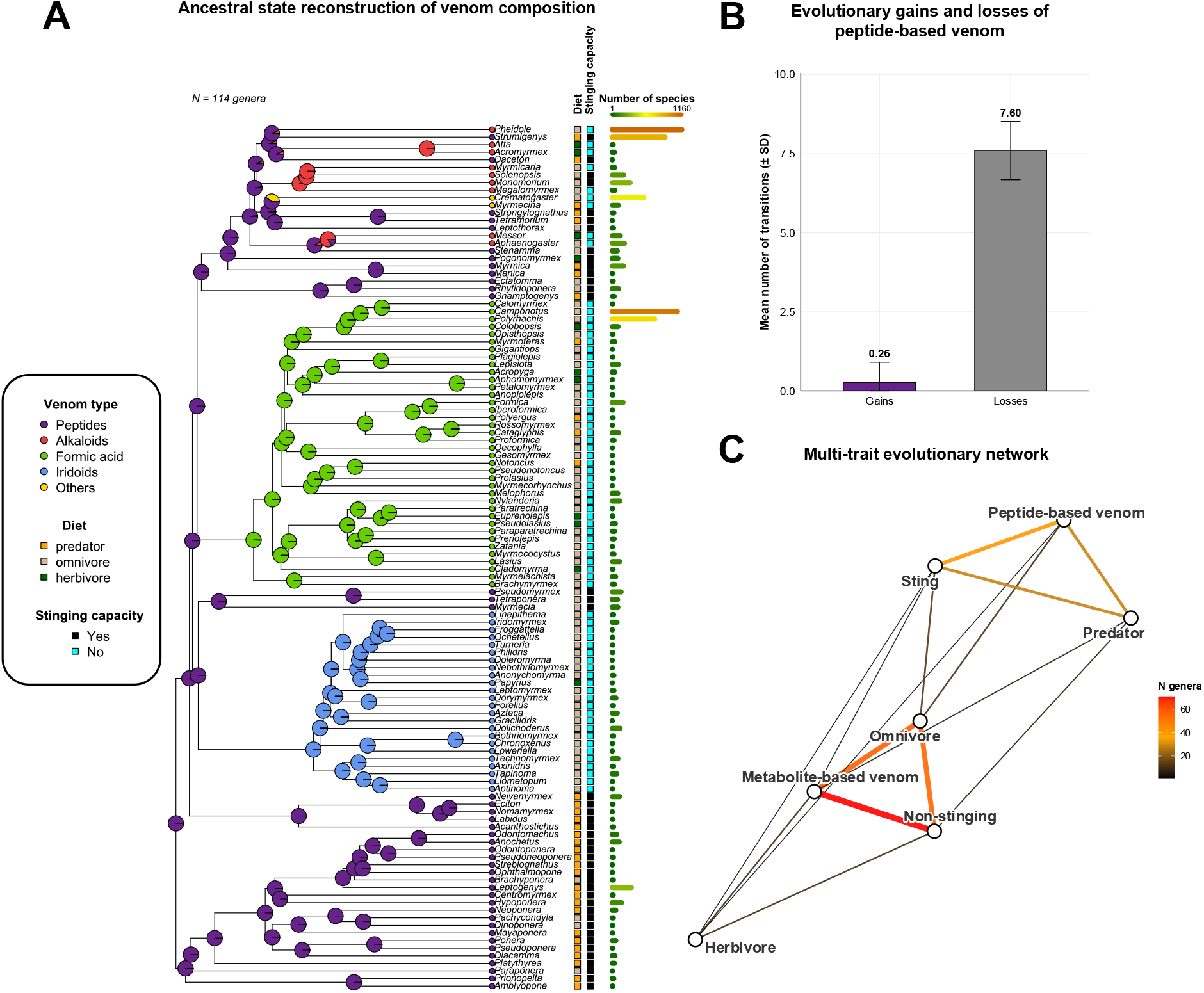
Evolutionary reconstruction of venom-type transitions and their association with sting capacity, diet and species richness across 114 ant genera. **(A)** Ancestral state reconstruction of venom chemical composition (pie charts, internal nodes; ace(), under the best-supported model selected by AIC and LRT from equal-rates (ER), symmetric (SYM) and all-rates-different (ARD) models). Tip circles show the observed venom type per genus.

Three Markov models of increasing complexity were compared to identify the most appropriate transition model for venom-type evolution before reconstructing ancestral states. The equal-rates (ER) model (AIC = 93.03) was favored over the symmetric (SYM) and all-rates-different (ARD) models (AIC = 97.24 and 115.10, respectively). A likelihood-ratio test confirmed that there was no significant improvement in fit under the ARD model (LRT = 15.93, df = 19, *P* = 0.662). Therefore, the ER model was used in all subsequent analyses. Stochastic character mapping was performed under this best-fit model to estimate the direction and frequency of evolutionary shifts in venom chemistry. Peptide-based venom exhibited asymmetric turnover, with an average of 0.26 gains versus 7.6 losses across 100 simulations (**Figure 2B**). Transitions occurred almost exclusively away from peptides (e.g., alkaloids: 3.97; others: 1.56; formic acid: 1.04; iridoids: 1.03), with negligible reverse transitions (≤0.21). This indicates that peptide-based venom was lost repeatedly and irreversibly over the course of diversification. To test whether the capacity to sting (Blanchard & Moreau, 2017) and diet (Greer & Moreau, 2021) are randomly distributed across the phylogeny or instead cluster among closely related genera, the phylogenetic signal was quantified using the D-statistic. Both the capacity to sting (D = −2.07) and predator diet (D = −1.07) showed phylogenetic clustering that exceeded Brownian expectation (*P* < 0.001 against a random distribution for both). Pagel’s lambda for the capacity to sting, reported for reference only, was 0.9999 (*P* = 5.43 × 10^−38^).

To assess whether venom type has evolved jointly with sting capacity and diet, correlated-evolution models were fitted for each trait pair (**Figure 2C**). Peptide-based venom and the ability to sting were strongly associated (41 out of 41 peptide-venom genera can sting versus only two out of 73 metabolite-based venom genera). The dependent (ARD) model fit better than the independent (ER) model (AIC = 87.03 versus 117.29, ΔAIC = 30.26, LRT = 36.26, df = 3, *P* = 6.61 × 10^−8^), as confirmed by Pagel’s test (likelihood ratio = 53.20, *P* = 7.73 × 10^−11^). Likewise, metabolite-based venom was significantly correlated with a dietary shift away from predation and toward omnivorous or herbivorous diets. This correlation was based on the joint distribution of venom type and diet across 114 genera: 68 metabolite-based/non-predator, 5 metabolite-based/predator, 12 peptide-based/non-predator, and 29 peptide-based/predator. A model in which venom type and diet evolve together (dependent ARD) fit the data far better than a model in which they evolve independently (ER) (AIC = 150.08 vs. 168.40; ΔAIC = 18.32; likelihood-ratio test: LRT = 32.32, df = 7, *P* = 3.54 × 10^−5^). This result was confirmed by Pagel’s test of correlated evolution (likelihood ratio = 22.06, *P* = 1.95 × 10^−4^).

Adjacent columns indicate diet and stinging capacity. Outer bars depict genus-level species richness N = number of genera analysed. **(B)** Mean number of evolutionary gains versus losses of peptide-based venom, estimated by stochastic character mapping (100 simulations under the best-fit model with the root fixed to ‘Peptide-based venom’). Gains correspond to transitions into a peptide-rich state, while losses correspond to transitions away from peptides towards alkaloid-, formic acid-, iridoid-or other metabolite-based venom. **(C)** A network showing the associations between venom type (peptide-based vs metabolite-based), stinging capacity (sting vs non-sting) and diet (predator, omnivore or herbivore). Edge width and colour indicate the number of genera sharing each trait combination (‘N genera’). Edges link venom type–diet, venom type–sting capacity and sting capacity–diet.

## IV. Why do ants repeatedly lose peptide-rich venoms?

With rare exceptions such as tetrodotoxin and anabaseine in nemertean worms (Williams, 2010; Göransson *et al*., 2019), animal venoms are dominated by peptides and proteins (Casewell *et al*., 2013). Although several ant lineages possess peptide-rich venoms delivered through a functional sting, ants diverge from the typical venom paradigm more than any other venomous animal group studied to date. Our review reveals at least seven independent transitions from ancestral peptide-based venoms to non-peptidic systems involving diverse alkaloids, formic acid, iridoids, esters, and other small organic compounds (**Section II; Figure 1**). These transitions are often linked to morphological modifications of the sting apparatus, including reduction or spatulate forms, as well as major shifts in venom deployment, such as topical application or spraying. Ancestral state reconstruction and stochastic character mapping confirm that peptide-based venom represents the ancestral condition in ants and has been lost repeatedly and has rarely, if ever, reversed over the course of diversification (**Section III; Figure 2A, 2B**).

Recent comparative analyses of ant sting morphology (Casadei-Ferreira, Richter & Economo, 2026) offer an ecological framework through which to interpret these patterns. Transitions from piercing to non-piercing sting systems are correlated with dietary diversification and changes in foraging strategies. These transitions reflect evolutionary trade-offs among predation efficiency, defense, and ecological flexibility. Piercing stings favor the immobilization of prey and ecological specialization, whereas non-piercing systems are associated with behavioral versatility, collective foraging, and plant-based diets. Our comparative analyses support and extend this framework: venom type, sting capacity and diet show significant correlated evolution across the ant phylogeny (**Section III; Figure 2C**), and transitions towards metabolite-based venom composition often parallel morphological modification of the delivery system (**Section III; Figure 2A**). Notably, most ant lineages that have evolved metabolite-based venoms belong to some of the most ecologically dominant, species-rich, and evolutionarily successful clades or genera (e.g., Formicinae, *Pheidole*, *Solenopsis*, *Crematogaster* and the fungus-growing ants). This potentially indicates that these transitions represent evolutionary innovations rather than simple losses. Often omnivorous or depending on plant-derived resources, these ants rely on collective foraging, numerical dominance, cooperative prey handling, and alternative defensive strategies, such as thickened cuticles, thoracic spines, or thanatosis (Blanchard & Moreau, 2017). Such traits likely relax selection on fast-acting, highly specific, injectable peptide toxins. These lineages tend to have venom dominated by multifunctional metabolites that combine insecticidal, defensive, antimicrobial, and communicative (alarm, recruitment, or trail) roles (**Section II**). This versatility likely facilitated their co-option across diverse ecological contexts.

Peptide toxins are presumed to be metabolically costly to biosynthesize since each molecule must be individually transcribed and translated (ribosomes, ATP/GTP), and often undergoes extensive post-translational modification (Lynch & Marinov, 2015). Small metabolites, by contrast, are typically generated by a limited set of enzymes, each of which can catalyse many turnovers over its lifetime (Bar-Even *et al*., 2011), making them comparatively less costly to produce, although enzymes of secondary metabolism tend to be less efficient than those of central metabolism (Bar-Even *et al*., 2011). This shift towards catalytically amplified, non-peptidic toxins may relax the energetic constraints on venom production, freeing resources for ecological expansion and diversification into novel niches.

This shift may also reflect nitrogen availability rather than energy alone. Predatory lineages obtain nitrogen directly and abundantly from animal prey, providing them with the resources to produce nitrogen-rich peptide toxins, whereas many lineages that have abandoned peptide venom rely instead on nitrogen-poor, carbon-rich resources such as honeydew, extrafloral nectar and fungal biomass, a well-documented driver of nitrogen limitation in ants (Davidson *et al*., 2003). Arboreal, canopy-dwelling lineages such as *Crematogaster* and the Dolichoderinae are particularly affected by this constraint (Davidson *et al*., 2003). Some other lineages have evolved symbiotic solutions to compensate for this limitation, including the nitrogen-recycling endosymbiont *Blochmannia* in *Camponotus* (Feldhaar *et al*., 2007), the conserved nitrogen-recycling gut microbiome of *Cephalotes* (Hu & Moreau, 2026), and nitrogen-fixing bacteria within fungus gardens (Pinto-Tomás *et al*., 2009), consistent with a broader association between herbivory and the acquisition of nitrogen-provisioning symbionts across the ant phylogeny (Russell *et al*., 2009). Under this hypothesis, metabolite-based venoms may be favored in nitrogen-limited lineages because they divert little or no nitrogen away from growth and reproduction, an advantage that is complete for nitrogen-free compounds such as formic acid or acetate esters, and more partial for alkaloids, which typically retain only one or two recycled nitrogen atoms per molecule rather than the dozens present in a peptide toxin.

In contrast, ant lineages possessing peptide-rich venoms have typically evolved under selective pressures associated with solitary predation, close-range antagonistic interactions, and vertebrate deterrence. In this context, peptide toxins, which are directly encoded by genes, offer clear advantages by forming adaptive venom cocktails. They can evolve rapidly through gene duplication, diversification, and neofunctionalization. The inherent evolvability of peptides under prey–predator arms races enables fine molecular targeting and high potency at low doses. These properties allow ants to continuously adjust their venom composition for offensive and defensive purposes, maintaining a competitive advantage in highly specialized or ecologically demanding niches.

The repeated emergence of metabolite-based venoms in ants raises fundamental questions about their evolutionary origins. Potential mechanisms may include *de novo* metabolic innovations, as exemplified by formic acid biosynthesis in Formicinae, where the involvement of the serine/glycine and THF metabolism pathway is well supported and relevant enzymatic activities have been identified in the venom gland, although the mechanisms underlying the accumulation of such high concentrations of formic acid in the venom reservoir remain unknown (see review by Koch *et al*., 2025). Mechanisms relying on endogenous ant metabolism are further supported by the identification of geraniol 8-hydroxylase activity in Dolichoderinae, which is key to iridoid biosynthesis (Datta *et al*., 2026). Alternatively, microbial symbioses, as documented for other toxin systems such as tetrodotoxin, may contribute essential precursors or metabolic steps, particularly when nitrogen must be incorporated despite nitrogen-poor diets (Feldhaar *et al*., 2007; Chau, Kalaitzis & Neilan, 2011; Nißl *et al*., 2025). Finally, horizontal gene transfer (HGT), as demonstrated in the whitefly *Bemisia tabaci*, which acquired a plant-derived detoxification enzyme (Xia *et al*., 2021), and documented for toxin gene families in centipede venoms (De León *et al*., 2025), could provide additional biosynthetic capabilities. Although this mechanism has yet to be investigated in the context of ant venom evolution, HGT events are recurrent in ants (Rinke *et al*., 2026). Future multi-omics studies combined with enzyme functional assays will be necessary to fully comprehend those biochemical mechanisms to elucidate how ants have repeatedly reinvented their venom systems and to determine the extent to which these innovations have contributed to their exceptional ecological and evolutionary success.

## V. Conclusions

1. Compared to other venomous lineages, ants depart markedly from the peptide-dominated composition typical of animal venoms, having independently abandoned peptide-rich venom for metabolite-based chemistries at least seven times.
2. These losses appear to be irreversible and are closely associated with the loss of stinging capacity and a shift from predation to omnivory or herbivory. This pattern may reflect an ecologically driven syndrome, rather than independent coincidences.
3. This shift toward metabolite-based venom accompanies some of the most species-rich and ecologically dominant ant lineages, whose multifunctional and presumably cheaper venom chemistries may have freed energetic and nitrogen resources for collective foraging, cooperative defense, and ecological expansion.
4. Peptide-rich venom persists where solitary predation or vertebrate deterrence selects for the rapid evolvability and precision targeting that gene-encoded toxins provide.
5. Determining how metabolite-based venoms repeatedly arose and how this transition shaped ant diversification will require genomic, metabolomic, and functional data from a broader range of genera.

## VI. Materials and Methods

A genus-level ant phylogeny was matched to trait data (e.g., venom type, capacity to sting, diet, and species richness) collected in this review and from other studies (Blanchard & Moreau, 2017; Greer & Moreau, 2021) and databases (AntWiki, 2026; Bolton, 2026) by genus name. Unmatched genera were excluded, and the order of the tree and traits was verified before each analysis. Venom type was classified into five categories (peptides, alkaloids, formic acid, iridoids, and others), and “unknown” genera were excluded from venom analyses. Diet was recorded as predator versus non-predator for binary tests. The final dataset included 114 genera with complete venom, diet, and sting data. Transition rates among the five venom-type states were fitted with fitMk (phytools) under equal rates (ER), symmetric (SYM), and all-rates-different (ARD) models. These models were compared using the Akaike information criterion (AIC) and a likelihood-ratio test (LRT; ER nested within ARD). Stochastic character mapping (make.simmap, phytools; 100 simulations; root fixed to peptides; seed = 123) used the best-supported model to estimate the mean gains and losses of peptide venom. Ancestral states were reconstructed with ace() under the same model. The phylogenetic signal in the capacity to sting and predator diet (both binary) was tested using the D-statistic (phylo.d, caper package; 1,000 permutations; seed = 1). Pagel’s lambda was additionally reported for the capacity to sting for reference only. We tested for correlated evolution between (i) peptide venom and the capacity to sting and (ii) peptide venom and predator diet with corHMM (independent ER vs. dependent ARD models; rate.cat = 1; compared by AIC and LRT) and Pagel’s test of correlated binary evolution (fitPagel). For the corHMM models, the AIC was computed as 2k − 2·logLik, where k is the number of free rate parameters in each model’s rate index matrix. All analyses were performed in R version 4.5.2 (2025) using the packages ape (v5.8.1), dplyr (v1.1.4), phytools (v2.5.2), ggplot2 (v4.0.1), ggrepel (v0.9.6), phangorn (v2.12.1), cowplot (v1.2.0), svglite (v2.2.2), grid (v4.5.2), corHMM (v2.8), igraph (v2.2.1), ggraph (v2.2.2), caper (v1.0.4), and ggtree (v4.0.4).

## VII. Acknowledgements

We thank Andrea Yockey for proofreading the manuscript.

## VIII. Funding

This work was supported by the ANR (plANTs, ANR-26-CE44-4945-01; Labex CEBA, ANR-10-LABX-25-01) and by the European Regional Development Fund (FEDER Guyane 2021– 2027, SelecTox project, GUY005177). L.R. was supported by a doctoral fellowship from AIBSI (Université de Guyane, ANR-22-EXES-0005).

## IX. Author contributions

Conceptualization: A.T., C.B.W., C.S.M. Formal analysis: A.T. Supervision: A.T., C.B.W., J.O., A.B., V.C. Writing – original draft: E.V.-M., L.R., J.O., A.T., C.B.W., A.D.

## X. Competing interests

The authors declare no competing interests.

## XI. Data availability

The genus-level trait dataset (venom type, stinging capacity, diet and species richness) and the R scripts used for the comparative analyses are available from the corresponding authors upon reasonable request.

## Notes

### Competing Interest Statement

The authors have declared no competing interest.

